# multicrispr: gRNA design for prime editing and parallel targeting of thousands of targets

**DOI:** 10.1101/2020.04.15.042861

**Authors:** Aditya M Bhagwat, Johannes Graumann, Rene Wiegandt, Mette Bentsen, Carsten Kuenne, Jens Preussner, Thomas Braun, Mario Looso

## Abstract

Targeting the coding genome to introduce single nucleotide deletions/insertions via Crispr/Cas9 technology has become a standard procedure in recent years. It has quickly spawned a multitude of methods such as Prime Editing, Crispr/Cas9 assisted APEX proximity labeling of proteins, or homology directed repair (HDR), for which supporting bioinformatic tools are, however, lagging behind. New applications often require specific guide-RNA (gRNA) design functionality, and a generic gRNA design tool is critically missing. Here we review gRNA designer software and introduce multicrispr, an R based tool intended to design individual gRNAs as well as gRNA libraries targeting many genomic loci in parallel. The package is easy to use, detects, scores and filters gRNAs on both efficiency and specificity, visualizes and aggregates results per target or Crispr/Cas9 sequence, and finally returns both genomic ranges as well as sequences of preferred, off target-free gRNAs. In order to be generic, multicrispr defines and implements a **genomic arithmetics framework** as a basis for facile adaptation to techniques yet to arise. Its performance and new gRNA design concepts such as **target set specific filtering** for gRNA libraries render multicrispr the tool of choice when dealing with screening-like approaches.

## Background

Crispr/Cas9, first reported in 1987 (Ishino et al. 1987) and later realized to be a prokaryotic immune system (Bolotin et al. 2005; Mojica et al. 2005; Pourcel et al. 2005), has given rise to a versatile molecular tool kit for genome engineering and analysis (Gasiunas et al. 2012; Jinek et al. 2012; Cong et al. 2013). Molecularly, the system comprises two components (Fig 1A): the Cas9 enzyme, which originally cuts double stranded DNA, and a guide RNA (gRNA). The latter consists of a *scaffold* linked to a 20 nucleotide *spacer* (N20, N = A, C, G, or T). When followed by an NGG *protospacer adjacent motif* (PAM) on the opposite strand, it guides the Cas9 nuclease to a genomic region of sequence complementarity.

**Figure 1.**
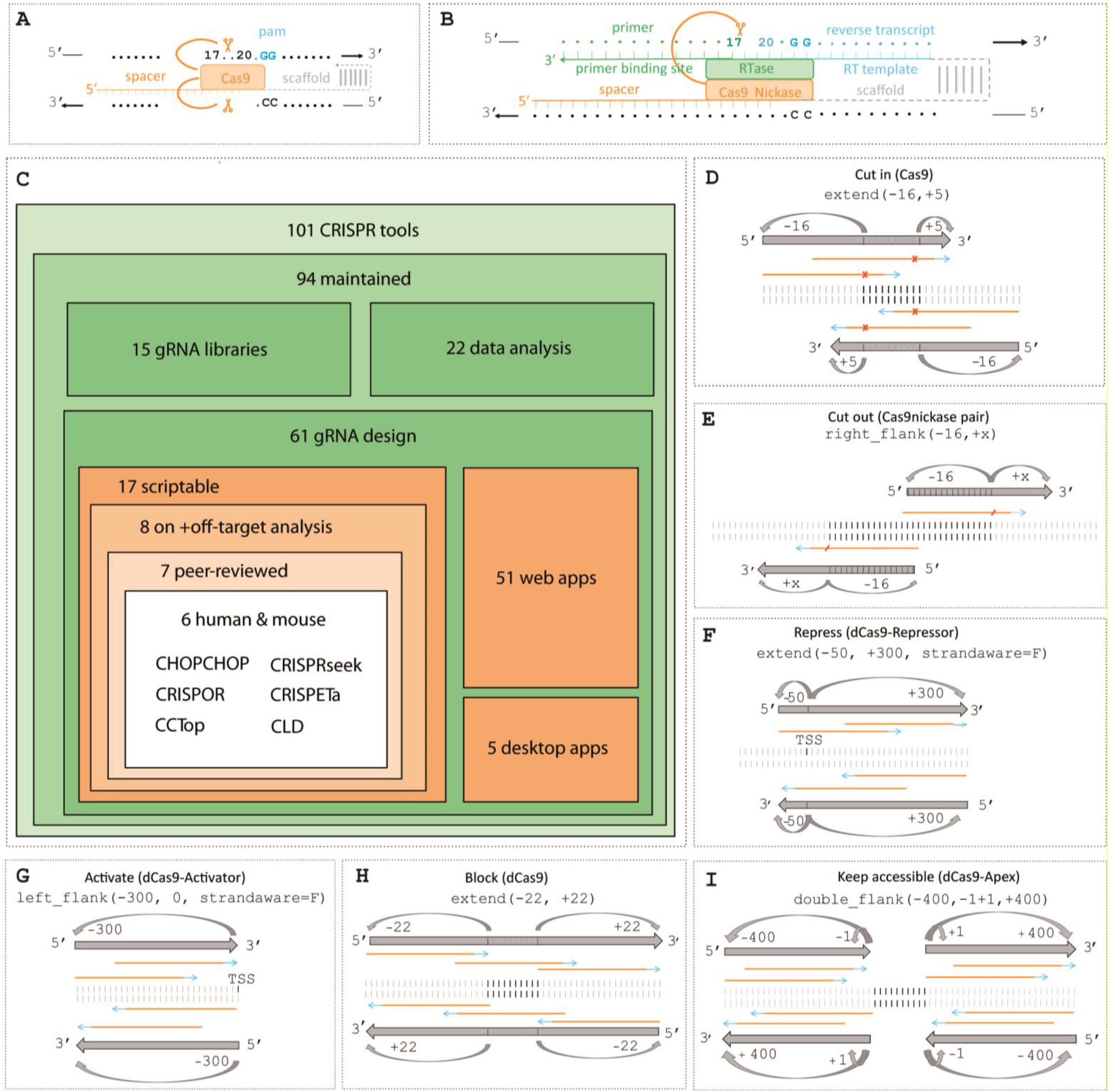
Schematic representation of Crispr/Cas9 application and arithmetics: **(A)** and **(B)** illustrate the basic Crispr/Cas9 mechanism and Prime Editing. Both systems target a genomic region based on complementarity to a 20-nucleotide spacer sequence (when followed by NGG on the opposite strand), and both involve cutting the pam-strand spacer after position 17 (double or single strand). The prime editor **(B)** additionally enables editing of sequence following nucleotide 17 through reverse transcription of a template (light blue, provided as a gRNA component), a process which is initiated through pairing of a primer binding site (another gRNA component) and primer (a portion of the spacer on the pam-strand). **(C)** A graphical overview of existing Crispr/Cas9 gRNA design tools as provided by (Torres-Perez et al. 2019) and their classification. **(D-I)** genomic arithmetics as needed for individual Crispr/Cas9 applications as indicated. Black lines represent the target range, orange arrows indicate the spacer sequences, blue arrows are pam sequences, orange crosses depict Cas9 cut sites, and large arrows mark the search region for spacer-pam sequences.

The Crispr/Cas9 application portfolio is constantly growing at considerable speed. One notable recent innovation is Prime Editing (Anzalone et al. 2019) (Figure 1B). This technology fuses a *Cas9 nickase* (cutting a DNA single strand) to a *Reverse Transcriptase*, and combines it with an extended guide RNA consisting of a spacer, a *primer binding site*, and a *reverse transcription template*. While the *spacer* continues to guide the complex to a genomic locus, the *primer binding site* also binds a region in the target DNA that serves as a primer for reverse transcription utilizing the additionally provided *reverse transcription template* (Fig 1B). This prime editor allows re-writing of up to 48 nucleotides at a specific locus of interest, enabling knockout, knockin, and precision editing.

A further example for a recently emerged Crispr/Cas9 application is parallel targeting of many loci with gRNA libraries, required for instance when targeting transcription factor binding sites (TFBS) or their neighborhoods (Shariati et al. 2019). Other screening oriented Crispr/Cas9-based applications include genome wide visualization (Zhou et al. 2017), or complex gRNA libraries to investigate cell fitness (Wegner et al. 2019). For such applications, the total number of gRNAs, or library complexity, directly correlates with effort and costs.

In general, several N20-NGG crispr sites may be identified for a genomic target region, but not all of them are equally suited. Thus, **gRNA design, defined as the process of finding an optimal gRNA**, involves two major tasks beyond the identification of sequences compatible with a target position: 1) off-target analysis to exclude spacers matching to other positions in the genome, while 2) on-target analysis involves selecting spacers that are expected to target the region of interest efficiently (using sequence-based prediction models).

Parallel targeting in a genome-wide context implies additional gRNA design needs. For one, the number of simultaneously targeted sequences may be large and processing efficiency thus is essential. In addition, e.g. when targeting TFBS sites, these target sequences are also prone to be very similar, as they conform to a consensus motif that can occur thousands of times. Consequently, Crispr/Cas9 spacers often match multiple targets. While this is undesirable for traditional Crispr-based techniques targeting single loci, it can be utilized for parallel targeting, allowing for a smaller gRNA set. The idea of parallel targeting thus requires differentiating between (mis)matches within the target set and to the genome, a process we define as **target set specific filtering** (TSSF).

To rise to the challenge of gRNA design, many software tools have been developed in recent years. As summarized in Fig 1C, the “WeReview:Crispr web table” (Torres-Perez et al. 2019) reports as many as 101 tools supporting CRISPR-based technology. When filtering for requirements essential to high throughput and reproducible results, such as current availability, support for gRNA design, access to a scripting interface, providing both on- and off-target analysis, being peer-reviewed, and supporting at least human and mouse as target organisms, this number reduces to only 6. Ordered by number of citations, these six are: CHOPCHOP ((Montague et al. 2014; Labun et al. 2016; Labun et al. 2019), 877 citations), CRISPOR ((Haeussler et al. 2016a; Concordet and Haeussler 2018), 532 citations), CCTop ((Stemmer et al. 2015), 329 citations), CRISPRseek ((Zhu et al. 2014), 118 citations), CLD ((Heigwer et al. 2016), 36 citations), and CRISPETa ((Pulido-Quetglas et al. 2017), 30 citations) (citations retrieved from google scholar on 17 February 2019).

As of March 2020, however, these tools largely do not provide functionality in support of more recently introduced applications of Crispr/Cas9. Of the six tools mentioned above, Prime Editing is currently supported by CRISPRseek only. Others are suffering from performance constraints (see Results/Benchmarking), limiting their usability when it comes to approaches requiring design of large gRNA libraries.

Existing tools aim to keep up with the ever growing pool of Crispr applications by continuous adaptation of their implementation and the occasional release of new versions including new functionality (Hanna and Doench 2020). Some miss a generalized framework and coding paradigm that allows for the efficient extension and adaptation to new applications while maintaining backward compatibility, and a continuous integration software design circuit. In this context, we propose **generic genomic arithmetics** as an essential feature to render a gRNA design tool suitable for the plethora of already established, as well as future Crispr-based strategies. For example, cutting within a target site (Fig 1D) requires a strand-specific [−16 +5] extension before searching for N20-NGG spacer-pam sequences, to ensure that Cas9, which has a cut site after nucleotide 17, cleaves within the target range. Excising a target site (Fig 1E) with a Cas9/Nickase pair, and possibly fixing it with homology directed repair (HDR), requires right flanking by [−16, +x] to find a Cas9/nickase pair in “PAM-out orientation” for excision (a Cas9/nickase pair in the [-x, +5] left flanks with a “PAM-in” orientation has been experimentally shown to be ineffective (Gearing 2018)).

These operations are strand-sensitive with reference to the coding sequence. Another example is the reduction of gene expression via a dCas9-repressor approach (Fig 1F), where a [−50, +300] extension of the TSS is required prior to spacer-pam sequence search (Gilbert et al. 2014). In contrast, when activating a gene with a dCas9-activator approach (Fig 1G), a [−300, 0] left-flanking of the transcription start site is required prior to spacer/pam sequence search (Doench 2020), allowing activator binding in relevant promoter/enhancer regions. Finally, blocking a target site (Fig 1H) requires a [−22, +22] target extension prior to spacer/pam search, ensuring that at least one nucleotide of the target area gets blocked. As indicated in (Fig 1I), targeting the vicinity of a sequence requires searching for crispr sequences in both flanks around the target site. The latter is for example needed for nucleoprotein complex purification via an affinity-tagged dCas9 (Liu et al. 2017), as well as Apex2-based protein interactor biotinylation (Myers et al. 2018). A subset of the existing software tools offer limited genome arithmetic functionality, such as options to specify a TSS offset between a target sequence and crispr spacer/pam sequences (e.g. CHOPCHOP, CRISPRSeek). However, the increasing variety of modern Crispr applications renders the flexibility resulting from genome arithmetics indispensable. Summarizing, an easily extensible Crispr design tool able to encompass both, current and future Crispr methods, will profit from a comprehensive genome arithmetics vocabulary that does not yet exist in this form.

## Results

### The multicrispr package

Motivated by the need for a gRNA design tool with generic functionality supporting a multitude of Crispr approaches, we developed the R package multicrispr. It has been designed to be highly performant as well as user-friendly, and provides comprehensive genome arithmetics vocabulary. As outlined in Fig 2A, a typical multicrispr workflow consists of five sequential steps with parametrization individual to specific Crispr applications. Each step provides optional plotting functionality, as exemplarily showcased for two applications in the next section. In order to integrate seamlessly into the R environment and the R/Bioconductor universe, a GRanges object (a core Bioconductor class) is returned as a final result, including information on both off- and on-target analysis. In addition, multicrispr generates human-readable and machine-parsable CSV files for further downstream processing.

**Figure 2.**
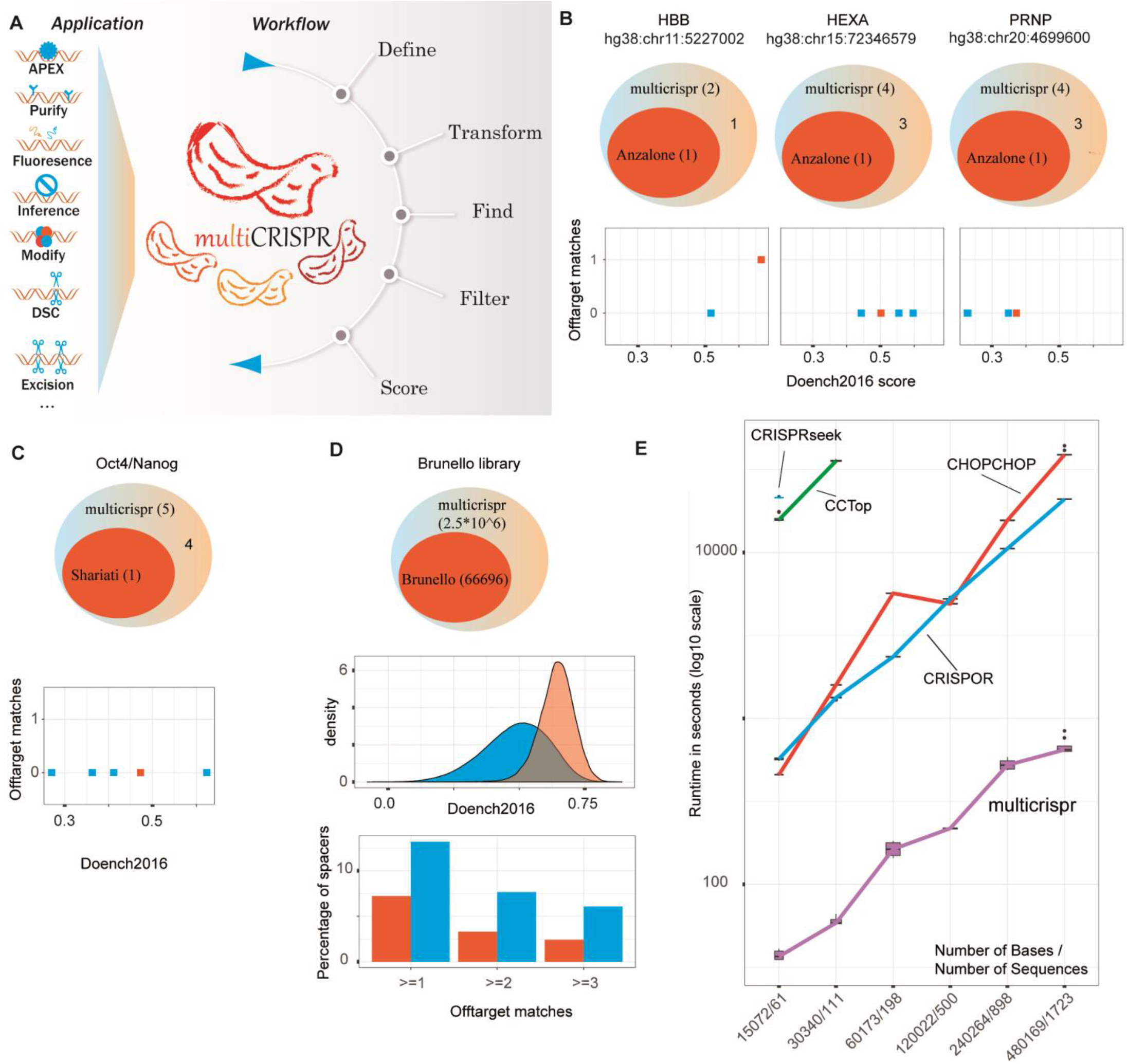
multicrispr workflow and validation: **(A)** Selection of supported crispr applications and workflow of multicrispr. **(B)** Overlap of Prime Editing spacer output of multicrispr and spacers used for the HBB sickle cell, HEXA Tay-Sachs, and PRNP Prion Disease locus, as given by (Anzalone et al. 2019). Scatter plots indicate scores and #mismatches given for all spacers found by multicrispr for the respective loci. **(C)** Overlap of multicrispr spacers and spacers used to block Oct4 TFBS [−151, −137] upstream of the Nanog gene, as utilized in (Shariati et al. 2019). Scatter plots indicate scores and #mismatches given for all spacers found by multicrispr for the respective loci. **(D)** Overlap of spacers identified with multicrispr for all Brunello exons (Doench et al. 2016). Density plot indicates scores for spacers specific for multicrispr (blue) and overlapping Brunello (red). Bar plots indicate # mismatches for these spacer sets as well. **(E)** Runtime comparison of gRNA design tools: the X-axis depicts increasing numbers of input sequences, while the Y-axis shows the total time needed by individual tools to design respective gRNAs on a log10 scale in seconds. Colors represent individual tools. Box plots represent repetitive processing of each input file (n = 10) to control for variability in computing performance.

The workflow starts with defining targets for genome engineering by either providing genomic coordinates or genomic identifiers. These targets are transformed into spacer/pam targets through *extension* and/or *flanking* (upstream, downstream) operations as required for the individual Crispr application performed (discussed above and in Fig 1). In the subsequent step, the transformed target ranges are searched for spacer/pam sequences. By default the wildtype S. pyogenes N20-NGG (spacer/pam) sequence is sought, but alternative spacer/pam sequences may be specified as well. The identified spacers are next filtered for specificity, resulting in spacers that match the targets, but no other genomic loci. The specificity function varies according to Crispr application. In the case of parallel targeting, exact matches with a variable amount of mismatches are allowed, and cross-target matches are considered on-target effects, as discussed earlier. For Prime Editing, which has been reported to be mismatch-free (Anzalone et al. 2019), only exact matches are considered and cross-target matches are rejected. Efficiency scores are added in a final step, providing extra guidance on which spacers to select for the experiment. Multicrispr supports the ‘Doench2014’ (Doench et al. 2014)) and ‘Doench2016’ (Doench et al. 2016) scoring models, the latter of which is currently the gold standard for gRNA efficiency prediction (Haeussler et al. 2016b).

### Validation

In order to validate the spacers identified by multicrispr, we applied our tool to gold standard targets in three different Crispr applications, with publications providing experimentally tested spacers of proven functionality. First we used multicrispr to identify spacers for Prime Editing the HBB sickle cell-, HEXA Tay-Sachs-, and PRNP prion disease loci, each of which was successfully prime edited by (Anzalone et al. 2019). Using multicrispr and as illustrated in Fig 2B, we confirmed spacers used by Anzalone et al. (2019) for all loci. Note that we were able to derive additional spacers targeting the same editing sites with scores and genomic mismatches comparable to the published controls.

In a second comparison, multicrispr was instructed to identify spacers blocking an Oct4 transcription factor binding site located upstream of the Nanog gene, reproducing work by (Shariati et al. 2019). In this case, multicrispr genome arithmetics functionality for upstream flanking was required to extend the target region by +/-22 nucleotides, as discussed earlier and detailed in Fig 1D-I. Aside from verifying the published spacer, multicrispr identified four additional spacers (Figure 2C), one of which is characterized by a higher Doench 2016 targeting efficiency score.

Finally, we used multicrispr to search for spacers in the exons targeted by the Brunello library (Doench et al. 2016), a validated gRNA library with 76,441 spacers targeting 19,114 transcripts (each transcript residing in a different gene, and each being targeted by up to four different gRNAs). After mapping to a current genome version and gene IDs (see methods), 66,696 of these spacers overlapped an exon, subsequently named as *Brunello exons*. Using these as target ranges, multicrispr identified more than 2.5 million spacers in total, including all 66,696 Brunello spacers (Fig 2D top). For each of them, the Doench2016 score was computed and a genome-wide (mis)match analysis was performed. As expected, the Brunello spacers are characterized by an enrichment in high Doench2016 scores (Fig 2D center). Genome (mis)match analysis revealed most identified spacers to be unique (Fig 2D bottom). Multicrispr, having been designed to scale to large datasets, performed both operations (Doench2016 scoring, genome (mis)match analysis) for the 2.5 million spacers within 1.1 hours and 1.6 hours respectively, utilizing a 15 Core / 128 GB RAM Linux virtual machine (see also Fig 2E and performance tests below).

Taken together, these examples confirm multicrispr’s ability to reproduce experimentally validated spacers efficiently for small-as well as large-scale screening-like applications.

### Benchmarking and feature comparison

From a practical point of view, performance of a computational tool is important, especially when it is not typically run on a horizontally scaling cluster, which is common for R analysis pipelines. In order to test the performance of multicrispr systematically, we performed comparative benchmarking with four other tools, involving spacer identification and on/off-target scoring for an iteratively increasing number of target sequences, utilizing the same 11 Core/132 GB RAM Linux machine. Random sequences fed to the tools were identical per run across the tools and an average of ten runs was calculated per target set size (Fig 2E). For CRISPRseek and CCTop, computation time for the two smaller exon sets were in the range of days to process and were thus excluded from further benchmarking. The remaining tools (CRISPOR, CHOPCHOP, and multicrispr) all met the challenge – with significant performance differences, however. While CHOPCHOP and CRISPOR needed processing time on the scale of hours, multicrispr finished the job within minutes (accelerating the search by a factor 21 when compared to the second fastest tool, CRISPOR). As a result, the Brunello exon example mentioned above is out of scale for all tested tools in terms of runtime.

Multicrispr provides another key functionality in the context of parallel targeting a large number of ranges. To the best of our knowledge, it is the only available tool that implements TSSF as introduced above, differentiating off-targets within a target set (which in parallel targeting experiments are considered on-targets) and others (“true” off-targets).

A detailed side by side comparison of gRNA design tools from the perspective of features implemented is shown in Table 1.

**Table 1:**
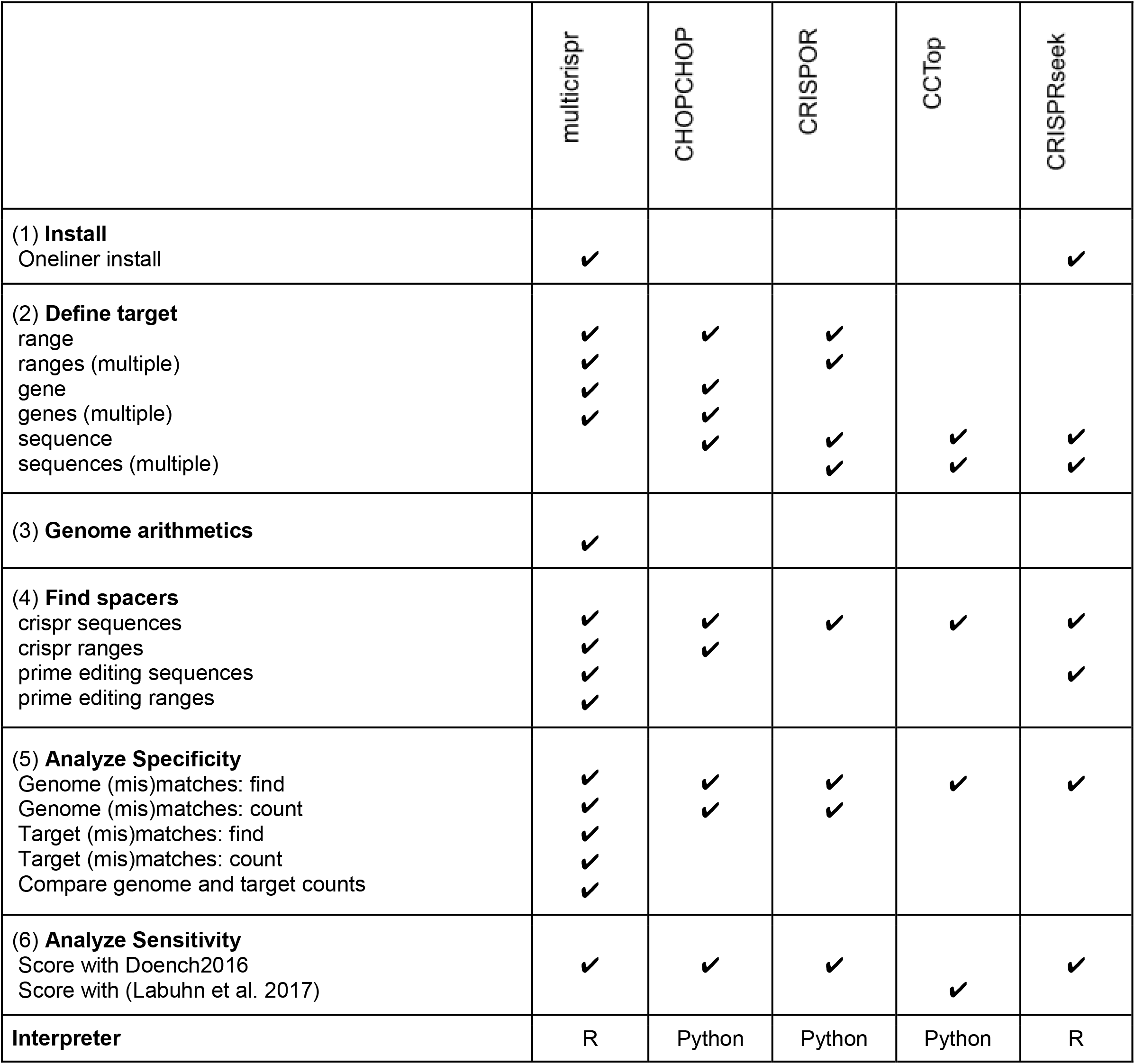
Feature comparison of gRNA design tools

In brief outline, multicrispr and CRISPRseek are both easy to install, while the others require the installation of long lists of dependencies. In terms of target definition, multicrispr processes targets defined as genomic ranges or gene names, instead of sequences. This allows to report the identified spacers in relation to their original target ranges and facilitates additional upstream or downstream analysis steps, such as the application of various BED format based tools. Although target range definitions are possible for CRISPOR and CHOPCHOP, the latter only accepts a single target per run. Explicit and extensive genome arithmetics functionality as defined in the background section is exclusively provided by multicrispr, with only limited implicit functionality provided by the other tools. Finding Prime Editing spacers (which imply additional sequence constraints), is only implemented by multicrispr and CRISPRseek. Specificity analysis is performed by all programs, however, the functionality provided is limited. Even though CHOPCHOP and CRISPOR also aggregate genome match counts, only multicrispr implements TSSF functionality, which additionally requires (mis)matching targets, counting target (mis)matches, and comparing target and genome mismatches. In comparison to CRISPRseek, the only other R package in the set, multicrispr facilitates the access to the Doench2016 scoring, building on the reticulate framework (Kevin Ushey 2019) and allowing within-R installation and single-line use of the python module azimuth from the Doench lab directly.

In summary we found multicrispr to lead other tools with respect to universality of application and extensibility, as well as smoothness of integration into its programming environment. Multicrispr thus defines a new standard with respect to performance and applicability in the context of large scale gRNA library design.

### Use Case 1: Prime Editing the sickle cell locus in the HBB gene

In order to exemplarily illustrate Prime Editing the sickle cell locus in the HBB gene on chr11 (see also Fig 2B), multicrispr must perform a bidirectional spacer search around the defined locus. The genome of interest as well as the genomic location of the SNP is thus specified as input (Fig 3A). Internally, the *find_pe_spacer* function call comprises a genome arithmetic extension ([−5, +nrt+16], nrt = number of reverse transcript nucleotides) that ensures the target site to be contained within the reverse transcript for editing (see Fig 1B for a discussion of prime editor architecture). In their seminal Prime Editing publication, Anzalone et al (2019) suggest nrt values up to 48, which is the value used here. Thus parameterized, multicrispr identifies six spacer-pam sites, two of them on the reverse, and four on the forward strand (Fig 3A). Subsequent evaluation by the filtering module finds four forward spacers to be target-specific, implying no interfering homology to other genomic sequences. In the next step, the scoring module assigned two spacers a high targeting efficiency according to Doench2016. Finally, the identified spacers are returned as a GRanges object with sequences for spacer, pam, primer, reverse transcript, as well as the 3’ extension.

**Figure 3.**
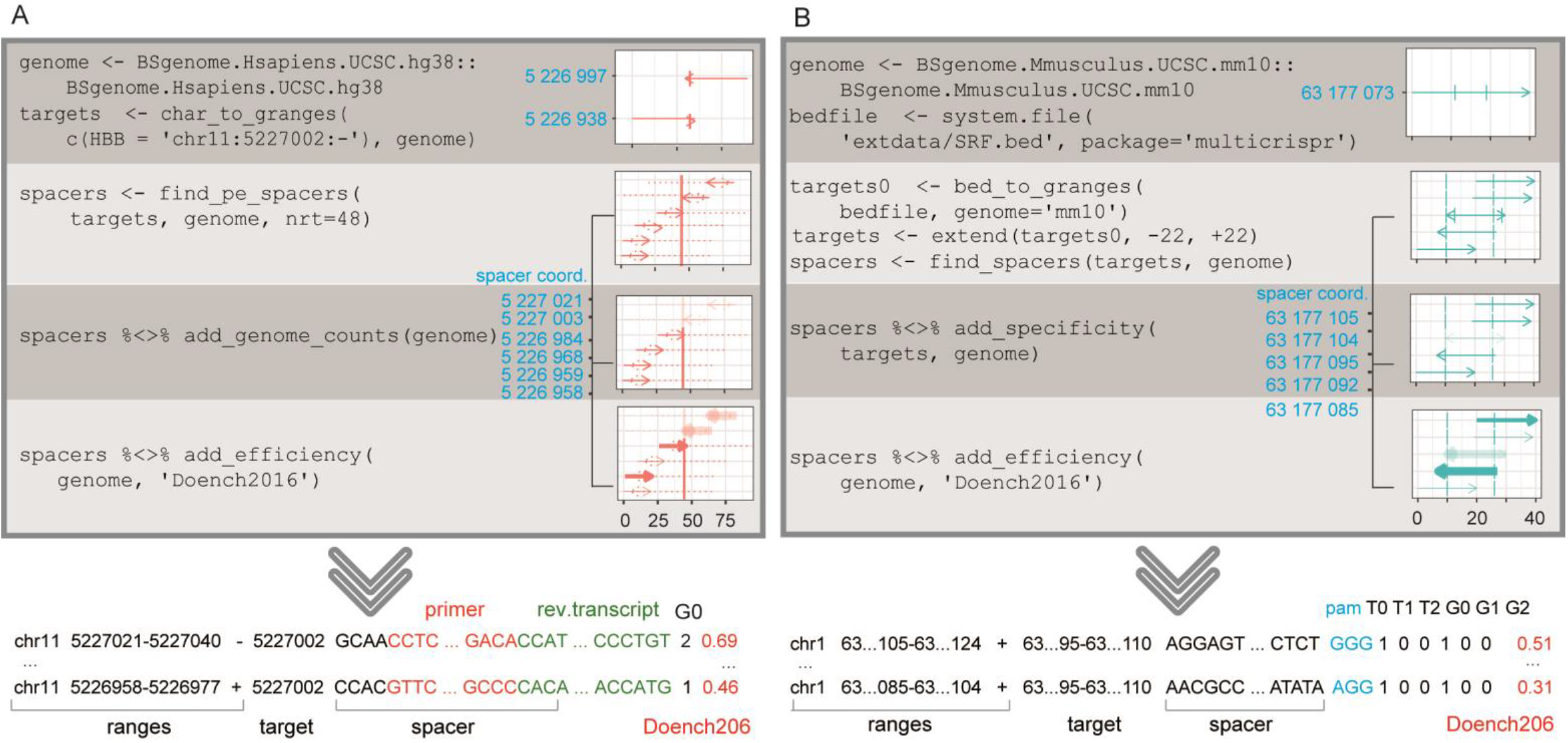
Use cases of multicrispr: **(A)** multicrispr code flow and visual output provided from deriving gRNAs for Prime Editing of the sickle cell locus in the HBB gene. Boxes indicate genomic locus on Y axis and range width on X. Primer binding site and reverse transcription template, jointly referred to as 3’ extension, are shown with dotted lines. Non-specific spacers are faded out (third box), targeting efficiency according to Doench2016 is mapped to line thickness. **(B)** multicrispr code flow and visual output provided from deriving gRNAs for the parallel targeting of 1,974 SRF binding sites. Boxes show one (of the 163) chr1 binding sites, with the start position mapped to Y and 1+width to X axis. Multicrispr finds 5 spacer-pam sites (of which 1 is a palindromic pair) for the blue binding site. Four of them are target-specific, two of which are predicted to have a good targeting efficiency. The resulting GRanges object is presented as a table (T0 = number of exact matches among targets, G1 = number of single nucleotide mismatches in the genome, etc.).

### Use Case 2: parallel targeting of transcription factor binding sites

In a second example, we utilize multicrispr’s functionality to derive gRNAs for a parallel targeting application involving the blocking of 1,974 binding sites of a transcription factor (here SRF) by a CRISPRi approach (Qi et al. 2013). On average, the SRF motif is sixteen nucleotides wide, and the corresponding sequences are provided to multicrispr as target ranges. As shown in Fig 3B, the ranges are first subjected to a [−22, +22] flank extension, to ensure that at least one spacer nucleotide is within the target range. The detection module of multicrispr subsequently derives strand-specific spacer/pam sites for the targets. Next, target set specific filtering is performed, excluding sequences with off-target (mis)matches, when they are outside of the target set. This is an essential step when dealing with large numbers of highly similar target ranges such as TF binding sites. For instance, two of the spacers shown in Fig 3b, while having a single exact match to the given target site, also have four off-target mismatches (three single, one double mismatch). However, all of these are located within the complete SRF target set, and consequently, are considered as on-target hits. Finally, multicrispr scoring functionality selects spacers with high efficiency potential, before a GRanges object including information on spacer (and target) ranges, spacer (and pam) sequences, on-target scores (Doench2016), and target and genome (mis)matches is returned.

## Discussion

Crispr/Cas9 is an increasingly versatile tool for genome engineering, with a high innovation rate regarding new applications. In general, the very first task in conducting a successful Crispr based experiment is a proper guide RNA design, choosing efficient gRNA spacers with minimal off- and maximal on-target activity. While experimentally validated gRNA libraries such as the Brunello library (targeting exons of the human genome), exist for the coding genomes of many model organisms, non-model organisms and non-coding genomes are lacking such accessible resources. For these and other cases with the need for custom designed spacers, gRNA design tools have been developed. We review a subset of these suited for high throughput and reproducible research, and compare them to our new generic tool multicrispr. We found that the increasing number of Crispr/Cas9 based applications and the trend towards large scale screening require a new generation of gRNA design tools, able to process both custom defined targets as well as large numbers of them. We benchmarked all scriptable on/off target analysis performing gRNA tools from the Torres et al. Crispr/Cas9 review table (with the exception of two, which we did not manage to install) and found none of them to be able to handle the dimensions required when e.g. targeting close to all human genes as represented by the Brunello library (Doench et al. 2016). In contrast, our tool multicrispr, designed for performance and generic usage, scales well to very large datasets. For instance, on a set of 1,723 mouse exons, multicrispr completed an order of magnitude faster (17 minutes) than popular alternatives such as CHOPCHOP (10 hours) and CRISPOR (5 hours).

Mass-targeting with Crispr/Cas9 libraries has also created interesting niche applications such as the parallel targeting of an overlapping target set of related sequences distributed in the genome (e.g. TFBSs). Such applications intriguingly turn conventions upside down: off-targets are no longer always off-targets, if they occur in the target set, and may even be desirable, allowing the use of a smaller gRNA set. In this context, we defined the term TSSF and implemented it and all further functionality needed to provide high quality gRNAs for such applications in multicrispr.

Aside from performance and multi target related tasks, many novel applications require flexible genome arithmetics functionality prior to spacer search. While some applications necessitate target extension, others need flanking, inverse targeting, or explicit avoidance of targets. To be able to handle each of these cases, a flexible, intuitive genome arithmetics functionality is required. Multicrispr provides this functionality combined with an easy-to-use, and intuitive “grammar” (how to define ranges, flanks etc.) inspired by the tidyverse paradigm of functional programming (Wickham 2019). In addition, multicrispr is intended to support the process of gRNA design by visual output functionality that intuitively documents each individual analysis step.

Driven by its unique functionality and performance, multicrispr paves the way for future custom resources. Examples that come to mind, but are beyond the scope of this introduction of the tool, are the application to annotated molecule subclasses such as lncRNA or miRNAs in a genome wide context. Other examples include the application of Prime Editing functionality to all 70,000 ClinVar (Landrum et al. 2014) sites to investigate how many of them may per se be targeted/ altered. A further straightforward application may be the generation of a global resource of gRNA pairs for the excision of complete exons, by making use of the flanking functionality of multicrispr for annotated exons as a target set and effectively targeting the introns.

In summary, multicrispr defines new standards for gRNA design in terms of performance, modularity and universality. It supports a plethora of Crispr/Cas9 based applications, including recent developments with the need for TSSF functionality.

## Material and Methods

### Speed comparison of tools

For performance tests, we utilized exon sequences of chr1 of mouse mm10 assembly. Starting with 15,000 bp, seven sets of exon sequences were created randomly, each with approximately twice the total length of the previous one in base pairs. Resulting test sets were saved as FASTA- and BED-files, respectively.

Each tool was installed according to its documentation in identical conda (Anaconda Development Team 2016) testing environments. Test sets were processed ten times with each tool and resulting values were summarized to control for variances in computing performance. The tools were parameterized as indicated in Supp Table 1.

**Supp Table 1:** Parameter settings used for benchmarking

| Tool name | Parameters |
| --- | --- |
| CHOPCHOP | --fasta<br>-G mm10<br>-t WHOLE<br>-v 2<br>-scoringMethod DOENCH_2016<br>-BED -GenBank |
| CRISPOR | mm10 -mm=2 --effScores=fusi |
| CCTop | --totalMM 2 --coreMM 2 |
| CRISPRseek | findgRNAsWithREcutOnly = FALSE, findPairedgRNAOnly = FALSE,<br>format = "bed"<br>BSgenomeName =<br>BSgenome.Mmusculus.UCSC.mm10<br>chromToSearch = "all"<br>txdb = TxDb.Mmusculus.UCSC.mm10.knownGene<br>rule.set = "Root_RuleSet2_2016"<br>annotatePaired = FALSE<br>PAM.pattern = "NGG\$"<br>allowed.mismatch.PAM = 1<br>topN = 1000000,<br>annotateExon = FALSE<br>fetchSequence = FALSE<br>max.mismatch = 2 |
| multicrispr | mismatches = 2<br>genome = BSgenome.Mmusculus.UCSC.mm10<br>method = 'Doench2016' |

### Brunello library validation

The Brunello library (Doench et al. 2016) represents a validated set of gRNAs comprising 76,441 spacers targeting 19,114 transcripts (each transcript residing in a different gene and each being targeted by four different gRNAs). We downloaded the Brunello gRNA set descriptions from addgene (https://www.addgene.org/static/cms/filer_public/8b/4c/8b4c89d9-eac1-44b2-bb2f-8fea95672705/broadgpp-brunello-library-contents.txt), which provide RefSeq mRNA identifier, cutsite position, and spacer orientation for each gRNA in the set. After excluding the positive controls, RefSeq mRNA were mapped to chromosomes and (unique) strands using biomaRt (Ensembl 99, Homo sapiens). Next, we were able to extend cut sites to full spacer ranges for 75,232 spacers / 18,810 transcripts using a [−17, +2] extension for ‘+’ spacers and a [−16, +3] extension for ‘-’ spacers. Subsequently, we extended each spacer to the (first) smallest, fully enclosing exon (i.e. both spacer and pam are contained in the exon), using Ensembl 99 exon models, as provided through Bioconductor’s AnnotationHub record AH78783. This was successful for 66,696 Brunello spacers/exons in 18,800 transcripts. The resulting set was used for multicrispr validation. Supp Table 2 details the number of spacers/transcripts targeted as a result of the reconstruction sequence.

**Supp Table 2:** number of spacers/transcripts targeted as a result of subsequent reconstruction steps

|  | spacers | transcripts |
| --- | --- | --- |
| Brunello | 77,441 | 19,114 |
| Remove positive controls | 76,441 | 19,114 |
| Add (canonical) chromosome mappings | 75,463 | 18,869 |
| Add (unique) strand mappings | 75,244 | 18,813 |
| Extend cut sites to spacer ranges | 75,232 | 18,810 |
| Extend to smallest, fully enclosing exon | 66,696 | 18,800 |

## Availability

Gitlab: https://gitlab.gwdg.de/loosolab/software/multicrispr

Website tool and manual: https://loosolab.pages.gwdg.de/software/multicrispr/index.html

Pre-calculated indices: https://s3.mpi-bn.mpg.de/minio/data-multicrispr-2020/

Bioconductor: Under revision

## Installation

Multicrispr can be installed from within an R environment with:

~~~
        install.packages(‘remotes’)
        remotes::install_git(‘https://gitlab.gwdg.de/loosolab/software/multicrispr.git’,
                           repos = BiocManager::repositories())
~~~

The python package azimuth (for Doench2016 on-target scoring) can be installed from within R with:

~~~
        install.packages(‘reticulate’)
        reticulate::conda_create(‘azienv’, c(‘python=2.7’))
        reticulate::use_condaenv(‘azienv’)
        reticulate::py_install(c(‘azimuth’, ‘scikit-learn==0.17.1’), ‘azienv’, pip = TRUE)
~~~

A Bowtie-indexed BSgenome (required for offtarget analysis) can be created with index_genome as shown below. This function needs to be run only once for any particular BSgenome. For the frequently used cases, we created pre-built indices, which are downloaded automatically. Beside the 28 organisms for which Bioconductor provides BSgenomes, another set of 224 organisms in twobit format through their AnnotationHub interface (Morgan 2019) is available. These can be converted to a BSgenome (Pages 2020), and then analyzed with multicrispr.

~~~
        BiocManager::install(‘BSgenome.Mmusculus.UCSC.mm10’)
        BiocManager::install(‘BSgenome.Hsapiens.UCSC.hg38’)
        index_genome(BSgenome.Mmusculus.UCSC.mm10::BSgenome.Mmusculus.UCSC.mm10)
        index_genome(BSgenome.Hsapiens.UCSC.hg38::BSgenome.Hsapiens.UCSC.hg38)
~~~

## Visualization

Graphs were generated via R. Illustrations in the context of genomic positions are generated via the multicrispr plot_intervals function, which can also be called explicitly. It operates on a GRange objects of any length

~~~
        bsgenome <- BSgenome.Mmusculus.UCSC.mm10::BSgenome.Mmusculus.UCSC.mm10
        bedfile <- system.file(‘extdata/SRF.bed’, package = ‘multicrispr’)
        targets <- bed_to_granges(bedfile, ‘mm10’, plot = FALSE)
        plot_intervals(targets)
~~~

